# Robust three-dimensional expansion of human adult alveolar stem cells and SARS-CoV-2 infection

**DOI:** 10.1101/2020.07.10.194498

**Authors:** Jeonghwan Youk, Taewoo Kim, Kelly V. Evans, Young-Il Jeong, Yongsuk Hur, Seon Pyo Hong, Je Hyoung Kim, Kijong Yi, Su Yeon Kim, Kwon Joong Na, Thomas Bleazard, Ho Min Kim, Natasha Ivory, Krishnaa T. Mahbubani, Kourosh Saeb-Parsy, Young Tae Kim, Gou Young Koh, Byeong-Sun Choi, Young Seok Ju, Joo-Hyeon Lee

**Author notes:** These authors contributed equally (Jeonghwan Youk, Taewoo Kim, Kelly V. Evans, Young-Il Jeong, Yongsuk Hur, Seon Pyo Hong). **Correspondence to**, Young Seok Ju, Associate Professor, KAIST, Daejeon 34141, Republic of Korea, Joo-Hyeon Lee, Group Leader, Wellcome-MRC Cambridge Stem Cell Institute, University of Cambridge, Cambridge CB2 A0W, United Kingdom, Byeong-Sun Choi, Division head, Division of Viral Disease Research, Center for Infectious Disease Research, Korea Centers for Disease Control & Prevention, Cheongju, Republic of Korea, Gou Young Koh, Director, Center for Vascular Research, Institute of Basic Science, Daejeon 34126, Republic of Korea, Young Tae Kim, Professor, Department of Thoracic Surgery, Seoul National University Hospital, Seoul 03080, Republic of Korea.

## Abstract

Severe acute respiratory syndrome-coronavirus 2 (SARS-CoV-2), which is the cause of a present global pandemic, infects human lung alveolar cells (hACs). Characterising the pathogenesis is crucial for developing vaccines and therapeutics. However, the lack of models mirroring the cellular physiology and pathology of hACs limits the study. Here, we develop a feeder-free, long-term three-dimensional (3D) culture technique for human alveolar type 2 (hAT2) cells, and investigate infection response to SARS-CoV-2. By imaging-based analysis and single-cell transcriptome profiling, we reveal rapid viral replication and the increased expression of interferon-associated genes and pro-inflammatory genes in infected hAT2 cells, indicating robust endogenous innate immune response. Further tracing of viral mutations acquired during transmission identifies full infection of individual cells effectively from a single viral entry. Our study provides deep insights into the pathogenesis of SARS-CoV-2, and the application of long-term 3D hAT2 cultures as models for respiratory diseases.

## Main

Several members of the family Coronaviridae are transmitted from animals to humans and cause severe respiratory diseases in affected individuals^1^. These include the severe acute respiratory syndrome (SARS) and the Middle East respiratory syndrome (MERS) coronavirus. Currently, Coronavirus disease 2019 (COVID-19), caused by severe acute respiratory syndrome coronavirus 2 (SARS-CoV-2), is spreading globally^2^ and more than 11.8 million confirmed cases with ~542K deaths have been reported worldwide as of 7^th^ Jul 2020^3^. The lung alveoli are the main target for these emerging viruses^4^.

To develop strategies for efficient prevention, diagnosis, and treatment, the characteristics of new viruses, including mechanisms of cell entry and transmission, kinetics in replication and transcription, host reactions and genome evolution, should be accurately understood in target tissues. Although basic molecular mechanisms in SARS-CoV-2 infection have been identified^5–8^, most findings have been obtained from experiments using non-physiological cell lines^9^, model animals, such as transgenic mice expressing human angiotensin-converting enzyme 2 (ACE2)^10^, ferrets^11^ and golden hamsters^12^, or from observation in clinical cohorts^13^ and/or inference from in-silico computational methods^14–16^. As a consequence, we do not fully understand how SARS-CoV-2 affects human lung tissues in the physiological state.

Development of three-dimensional (3D) stem cell-derived organotypic culture models, conventionally called organoids, has enabled various physiologic and pathological studies using human-derived tissues *in vitro*^17–19^. Organoid models established from induced pluripotent stem cells (iPSCs), or adult stem cells in the human kidney, intestine, and airway have been used to investigate SARS-CoV-2 pathogenesis^20–23^. Although human alveolar type 2 cells (hereafter referred to as **hAT2s**) are believed to be the ultimate target cells for SARS-CoV-2, their infection model has not previously been introduced.

### Long-term 3D culturing of hAT2s without mesenchymal niche

We have established feeder-free, 3D hAT2 organoids (hereafter referred to as **hAOs;** definition of organoid is available at **ref. ^24^**) with defined factors which support molecular and functional identity of hAT2 cells over multiple passages, showing substantial improvements from the previous application of co-culture models^25–27^. Briefly, single-cell dissociated hAT2 cells were isolated by fluorescence-activated cell sorting (FACS) for the hAT2 surface marker HTII-280 (CD31^−^CD45^−^EpCAM^+^HTII-280^+^)^25,28^(**Fig. 1a**; **Extended Data Fig. 1a**). Isolated HTII-280^+^ cells showed higher expression of AT2 cell marker SFTPC, while HTII-280^−^ cells revealed higher expressions of basal cell marker TP63 and secretory cell marker SCGB1A1 (**Extended Data Fig. 1b**). We then plated HTII-280^+^ hAT2 cells into Matrigel for 3D cultures with our expansion medium, supplemented with CHIR99021, RSPO1 (R-spondin 1), FGF7, FGF10, EGF, NOG (Noggin), and SB431542, that are known to support the growth of human embryonic lung tip cells^29^. HTII-280^−^ cells were also cultured under conditions supporting human bronchial (airway) organoids (hereafter referred to as **hBO**s) that have previously been reported^30^. hAOs established from single hAT2 cells grew up to 4 weeks with heterogeneous morphology including budding-like and cystic-like structures consisting of mature AT2 cells expressing pro-SFTPC, HTII-280, and ABCA3, as well as exhibiting uptake of Lysotracker, a fluorescent dye that stains acidic organelles such as lamellar bodies^25^ (**Fig. 1b and 1c).** In contrast, hBOs grew quickly by day 14 with cystic-like structures consisting of a number of airway cell types, including KRT5^+^TP63^+^ basal cells and SCGB1A1^+^ secretory cells, as previously reported^30^ (**Fig. 1d and 1e**). WNT activation was identified as an essential factor for hAO formation, because no colony formation was found in the absence of WNT activator CHIR99021 in culture (**Extended Data Fig. 1c**). Importantly, our culture system allows the long-term expansion (>10 months) of hAT2s, although colony forming efficiency varied between tissue samples and reduced at later passages (**Extended Data Fig. 1d**). Over passaging via single cells, hAT2s consistently formed organoids, exhibiting SFTPC expression following 9 months of continuous cultures, although growth began to slow, as evident by reduced organoid size and lower forming efficiencies (**Extended Data Fig. 1d and 1e**). Alveolar type 1 cells (hAT1s) expressing HOPX and PDPN were also observed during early cultures, demonstrating differentiation capacity of hAT2 cells in our hAOs (**Extended Data Fig. 1f**), although later passages exhibited loss of these cells.

**Figure 1.**
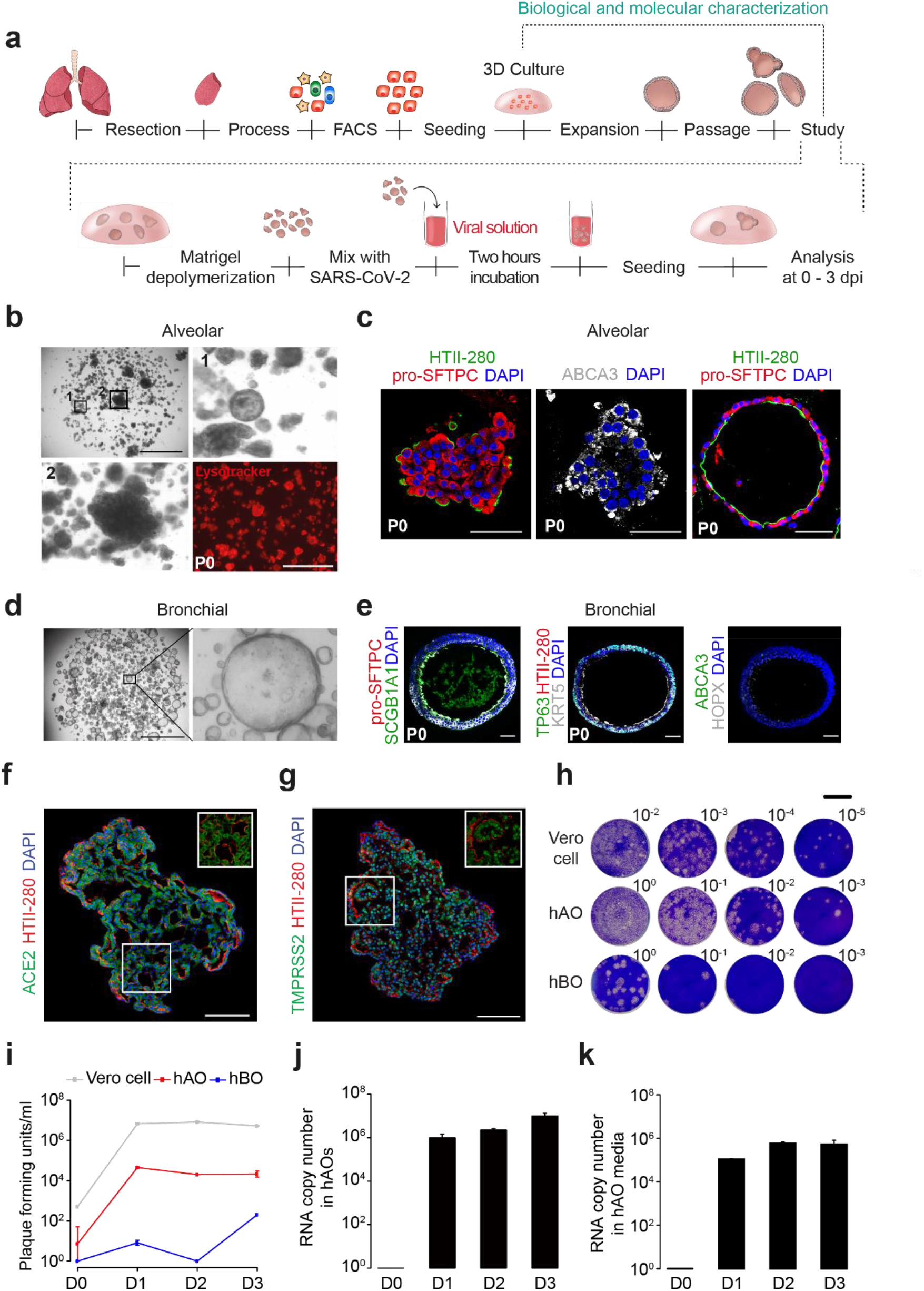
Establishment of long-term and three-dimensional human alveolar type 2 cell culture and SARS-CoV-2 infection in the three-dimensional model system. **a,** Schematic diagram outlining the method for dissociation and processing of human adult lung parenchymal tissues for *in vitro* three-dimensional cultures (top). Schematic illustration of SARS-CoV-2 infection experiments in this study (bottom). **b,** Representative bright-field images of hAOs derived from HTII-280^+^ hAT2 cells at day 28 in culture. Insets (top left) show high-power view of cystic-like (top right) and budding-like (bottom left) alveolar organoids. Scale bar, 2000 μm. Of note, the majority of hAOs exhibited uptake of Lysotracker (red), indicative of mature AT2 cells (bottom right). Scale bar, 1000 μm. **c,** Immunofluorescent staining of hAOs expressing AT2 markers. Left and right; HTII-280 (for hAT2, green), pro-SFTPC (for hAT2, red). Middle; ABCA3 (for hAT2, white). DAPI (blue). Scale bar, 50 μm. **d,** Representative bright-field images of hBOs derived from HTII-280^−^ non-hAT2 cells at day 14 in culture. Insets (left) show high-power view (right). Scale bar, 2000 μm. **e,** Immunofluorescent staining of hBOs expressing airway lineage markers. Left; SCGB1A1 (for secretory, green), pro-SFTPC (for hAT2, red). Middle; KRT5 (basal, white), TP63 (basal, green), HTII-280 (for hAT2, red). Right; ABCA3 (for hAT2, green), HOPX (for hAT1, white). DAPI (blue). Scale bar, 50 μm. **f,** Immunofluorescent staining of ACE2 (green) and HTII-280 (red) in hAOs. HTII-280 is stained in the apical membrane of hAT2 cells. Scale bar, 100 μm. **g,** Immunofluorescent staining of TMPRSS2 (green) in hAOs. Scale bar, 100 μm. **h,** Representative images for plaque assay at 3 dpi. Dilution factors are shown in numbers on the right upper corner. Scale bar, 1 cm. **i,** Plaque assay showing that SARS-CoV-2 actively replicates in hAOs at 1 dpi. SARS-CoV-2 can infect hBOs, but the viral amplification is much lower than in hAOs. Given the different culture and viral infection techniques between 2D Vero cells and 3D organoids, direct comparison is not applicable between Vero cells and hAOs. Error bars represent SEM. n=2. **j,** Quantitative PCR (qPCR) analysis for measuring the viral RNA levels in lysed hAOs. Error bars represent SEM. n=3. **k,** qPCR analysis for measuring the viral RNA levels in hAO media. Error bars represent SEM. n=3.

### Robust infection of SARS-CoV-2 into hAOs

The expressions of ACE2 and TMPRSS2, which are necessary for SARS-CoV-2 infection, were observed in the membrane and cytoplasm of hAO cells (**Fig. 1f and 1g; Extended Data Fig. 1g**). We next infected hAOs and hBOs with SARS-CoV-2 at a multiplicity of infection (**MOI**) of 1. The viral particles were prepared from a patient (known as **KCDC03**) who was diagnosed with COVID-19 on 26^th^ Jan, 2020, after traveling to Wuhan, China^31^. Vero cells were also infected as a positive control, although this was not directly comparable to our 3D models due to different technical procedures.

Infectious virus particles increased to significant titers in hAOs (**Fig. 1h-1k; Extended Data Fig. 2**), reaching maximum levels within the 1st day post infection (**dpi**), suggesting that full infection occurs within 1 day from viral entry to hAOs. In hBOs, the increment of viral particles was observed as consistent with another study^32^, but their titers were <100 times lower than hAOs (**Fig. 1h and 1i**; **Extended Data Fig. 2**). In line with viral particles, the amount of the viral RNA in hAOs and in its culture supernatant reached a plateau at 1 dpi (**Fig. 1j and 1k**). Although infected Vero cells exhibited significant cytopathic effects at 1 dpi, typically cell rounding, detachment, degeneration and syncytium formation^9^, SARS-CoV-2 infected hAOs and hBOs did not show prominent macroscopic pathologies up until 10 dpi.

### Structural changes of hAOs to SARS-CoV-2 infection

Immunostaining for double-stranded viral RNA (**dsRNA**) and nucleocapsid protein (**NP**) of SARS-CoV-2 identified widespread viral infection in hAT2 cells co-expressing pro-SFTPC and ACE2 in hAOs (**Fig. 2a and 2b; Extended Data Fig. 3**). To further determine subcellular events at a higher resolution, transmission electron microscopic analysis was performed at 2 dpi (**Fig. 2c-2l; Extended Data Fig. 4**). Most cells (~80%), containing secretory vesicles and surfactant proteins (lamellar bodies^25^), showed discernible viral particles in the cytoplasm. A fraction of cells in hAOs showed much higher viral burdens than other cells, with as many as 500 copies in the 100 nm section, implying that >10,000 SARS-CoV-2 particles per cell. Aggregated viral proteins, which appeared as electron-dense regions near nuclei^33^, were also detected (**Fig. 2c**). Accordingly, several key pathogenic phenotypes were observed in the infected hAOs. Alveolar cells with enormous vacuoles were frequently observed (**Fig. 2c, 2f, 2h, 2i**), similar to cytopathic signals in Zika virus^34^. Double-membrane vesicles (DMVs), subcellular structures known as viral replication sites frequently seen in the early phase of infection^35,36^, were observed in the vicinity of zippered endoplasmic reticulum in a small fraction of hAO cells (**Fig. 2j and 2k**). Viral particles were dispersed in the cytosol (**Fig. 2e**) or enclosed in the small vesicular structures (**Fig. 2f and 2i**). Diverse forms of viral secretion were also observed mainly through the apical surfaces of hAT2 cells (**Fig. 2g and 2l**). More ultrastructural pathologies are available at **Extended Data Fig. 4** and the EMPIAR data archive (see Data availability).

**Figure 2.**
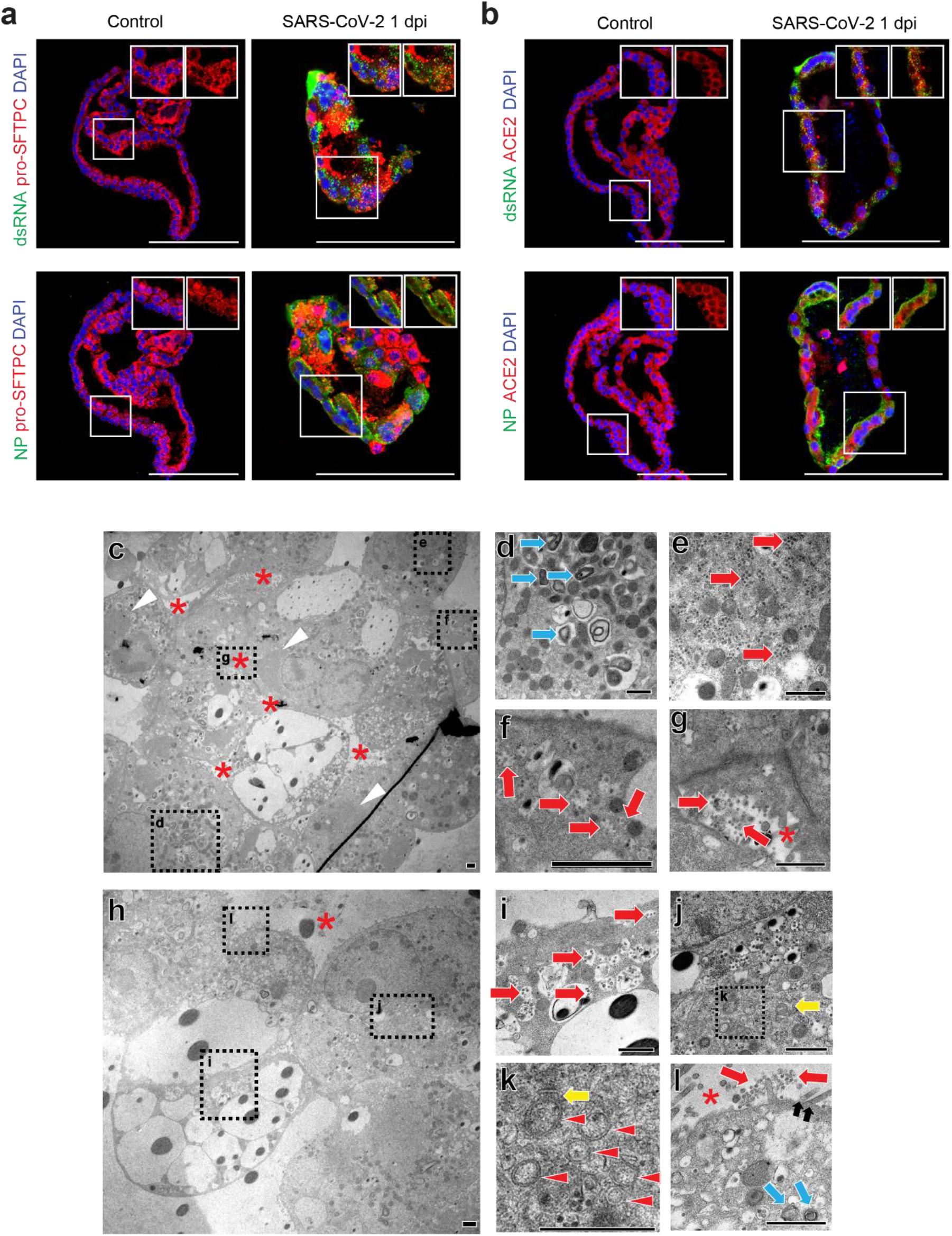
Confocal and transmission electron microscopic images of SARS-CoV-2 infected hAOs. **a,** SARS-CoV-2 infected hAOs at 1 dpi. Viral nucleocapsid protein (NP) and double-stranded RNA (dsRNA) are costained with pro-SFTPC. At 1 dpi, SARS-CoV-2 highly infects hAOs which show punctuated pattern of pro-SFTPC (red). Scale bar, 50 μm. **b,** The viruses are identified by dsRNA (top right) and NP (bottom right). Infected hAOs express ACE2 (red). Viral dsRNA (green) appears punctuated. More than 80% of hAT2s are infected at 1 dpi. Scale bar, 50 μm. **c,** Transmission electron microscopic image of an entire hAO structure. Small alveolar spaces are shown between hAT2s (red asterisk). Aggregated viral particles are sometimes observed in electron-dense areas of the hAOs (white arrowhead). Scale bar (c-l), 1 μm. **d,** Lamellar bodies (skyblue arrow), consisting of pulmonary surfactants, are frequently observed in hAOs. **e,** Hundreds of SARS-CoV-2 particles in the cytosol of hAT2 cells (red arrow). **f,** Multiple viral particles in vesicular structures. **g,** SARS-CoV-2 viral particles secreted into an alveolar space of hAOs. **h,** Another low magnification image of a hAO. Some cells contain large pathologic vacuoles. **i,** Large vacuoles usually do not include virus particles, but small vesicles contain many virus particles. **j,** Viral containing vesicles, aggregated in the vicinity of zippered endoplasmic reticulum (ER; yellow arrow). **k,** Double membrane vesicles (red arrowhead) located near zippered ER. **l,** Secreted viral particles in the apical lumen of hAOs. Microvilli (black arrow) and lamellar body (skyblue arrow) are shown at the apical side of hAT2s.

### Transcriptional changes of hAOs to SARS-CoV-2 infection

From strand-specific deep RNA-sequencing, we explored gene expression changes in the infected hAOs. Indeed, a set of human genes were differentially expressed as infection progressed (i.e., 0, 1 and 3 dpi), although most genes showed good correlations (**Extended Data Fig. 5a** and **Supplementary Table 1**). Cytokeratin genes (including *KRT16, KRT6A*, *KRT6B*, and *KRT6C*), genes involved in keratinization (including *SPRR1A*), cytoskeleton (including *S100A2*) and cell-cell adhesion genes (including *DSG3*), were significantly reduced to ~2-3% in hAOs at 3 dpi (**Fig 3a and 3b**). Many more genes were upregulated in the infected hAOs specifically at 3 dpi. In particular, transcription of a broad range of interferon-stimulated genes (**ISGs**), known to be typically activated by type I and III interferons^37^, were remarkably increased (**Fig. 3a and 3b**). These genes include interferon induced protein genes (such as *IFI6*, *IFI27*, *IFI44*, *IFI44L*), interferon induced transmembrane protein genes (such as *IFITM1*), interferon induced transmembrane proteins with tetratricopeptide repeats genes (*IFIT1*, *IFIT2*, *IFIT3*), 2’-5’-oligoadenylate synthetase genes (*OAS1*, *OAS2*), and miscellaneous genes known to be involved in innate cellular immunity (*MX1*, *MX2*, *RSAD2*, *ISG15*). These genes were expressed to >20 times higher levels in hAOs at 3 dpi than at 0 dpi. Many other known ISGs also showed moderate inductions (2-20 times) at 3 dpi, including *BTS2* (~15 times), *OAS3* (~11 times), *HERC5* (~15 times), *HERC6* (~12 times) and *USP18* (~12 times). Antiviral functions are known for these ISGs^38^ including (1) inhibition of virus entry (*MX* genes, *IFITM* genes), (2) inhibition of viral replication and translation (*IFIT* genes, *OAS* genes, *ISG15*, *HERC5*, *HERC6*, *USP18*) and (3) inhibition of viral egress (*RSAD2* and *BST2*). Of note, given that immune cells are absent in our culture system, the innate immune response was completely autologous to alveolar cells, mimicking the initial phase of SARS-CoV-2 alveolar infection. In line with the notion, innate induction of some type I and type III interferons was observed. Of the 20 interferon genes, an interferon beta gene (*IFNB1*) and three interferon lambda genes (*IFNL1, IFNL2*, and *IFNL3*) showed significant transcriptional induction, although their absolute changes were not substantial (**Fig. 3c**). The surface receptors of interferons were stably expressed in hAO cells without reference to viral infection (**Fig. 3c**). Downstream signalling genes of the receptors were also upregulated such as *STAT1* (~3.5 times), *STAT2* (~2.5 times) and their associated genes *IRF1* (~2.4 times) and *IRF9* (~6.4 times). Of note, *IRF1* is known to be specific to type I interferon responses^39^, while type I and type III ISGs are generally overlapping ^40^.

**Figure 3.**
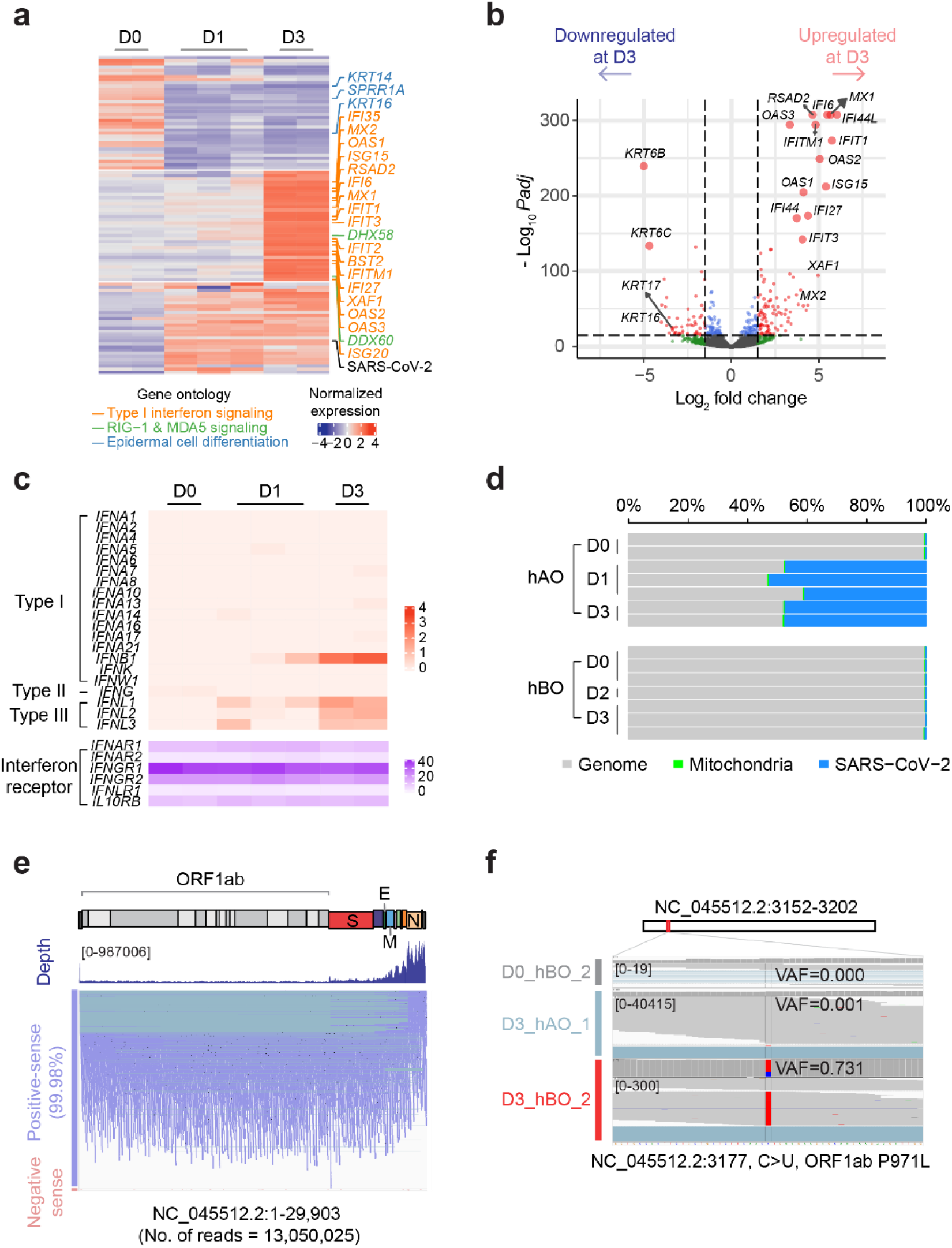
RNA-seq analyses of infected hAOs and hBOs. **a,** Heatmap of the most variable 100 genes among three groups of hAOs at 0, 1 and 3 dpi. Genes related to type I interferon signal pathway are highly elevated at 3 dpi, while SARS-CoV-2 expression reach plateau at 1 dpi. **b,** A volcano plot showing differentially expressed genes between hAOs at 0 dpi and 3 dpi. Most highly upregulated genes are in type I interferon signaling pathway. **c,** Transcriptional changes of interferon genes in the infected hAOs. The rise of *IFNB1, IFNL1, IFNL2, and IFNL3* are observed in hAOs. **d,** Proportion of viral RNA reads in the hAO and hBO transcriptomes. Compared to hAOs, the portion of viral reads is much lower in hBOs. **e,** Positive-sense viral RNAs are dominant (99.98%) over negative-sense viral RNAs. **f,** An example of missense mutation (NC_045512.2: 3,177C>U) detected from hBO transcriptome at 3 dpi. The variant allele fraction (VAF) of the substitution is 0.731 in an infected hBO sample, much higher than that in other infected cells.

In addition to ISGs, genes in the viral sensing pathway in cytosol showed increased expression in the infected hAOs at 3 dpi, for example, *DDX58* (official gene name of *RIG-1*, from 1.9 to 25.0 TPM), *IFIH1* (also known as *MDA5*, from 5.8 to 30.3 TPM), and *TLR3* (Toll-like receptor 3, from 1.3 to 3.7 TPM), *IRF7* (Interferon regulatory factor 7, from 4.4 to 29 TPM) and *IL6* (0.6 to 2.5 at 1 dpi; proinflammatory factor).

Notably, these transcriptional changes were much stronger in hAOs than in hBOs. In the similar transcriptome profiling of the infected hBOs, the genes aforementioned were not significantly altered (**Supplementary Table 2** and **Extended Data Fig. 5b**). In addition, we identified few hBO specific differentially expressed genes (**Extended Data Fig. 5c**). This finding implies cellular tropism of SARS-CoV-2 viral infection.

### Expression of SARS-CoV-2 genes in the infected organoids

We further analysed the viral RNA sequences obtained from the infected models. In agreement with the plaque assay (**Fig. 1h and 1i**), relative transcription of SARS-CoV-2 genes plateaued by 1 dpi (**Fig. 3d**), which is earlier than the host gene expression changes. Approximately 50% of the RNA sequencing reads were mappable to the SARS-CoV-2 genome in hAOs from 1 dpi (**Fig. 3d**), indicating prevailing viral gene expression in infected hAO cells as observed in Vero cells^6^. Of note, the proportion of viral transcripts was much lower in the infected hBOs.

Transcripts from SARS-CoV-2 were not mapped uniformly to the viral genome sequence, but 3’ genomic regions, where canonical subgenomic RNAs are located, showed much higher read-depth in all samples, consistent with the previous report^6^ (**Fig. 3e**). The vast majority of viral RNA sequences produced from the infected hAOs and hBOs was in the orientation of positive-sense RNA strands (**Fig. 3e**; for example, 99.98% vs 0.02% for positive- and negative-sense RNAs, respectively, from hAO at 1 dpi). This is in good agreement with the nature of SARS-CoV-2, which is an enveloped, non-segmented, and positive-sense RNA virus.

By cross-comparison of viral RNA sequences produced from a total of 11 infected hAOs (n=5) and hBCs (n=6), we identified 20 viral base substitutions (**Supplementary Table 3**). No mutation was at 100% variant allele fraction (**VAF**) and exclusive to an infected sample. Instead, sequence alterations showed a broad range of quasispecies heterogeneity in each culture (VAF ranges from 0.1% to 73.1%; **Fig. 3f**), and a large proportion of the mutations (n=16; 80%) were shared by two or more infected models (by the cut-off threshold of 0.1%). Therefore, we speculate that most of these sequence changes were originally present in the pool of viral particles before their inoculation. Given the fact that these viral particles were prepared from one of the earliest COVID-19 patients, our finding suggests that mutations can accumulate in the viral genomes in a small number of rounds of viral transmissions, and appear with dramatic changes in quasispecies abundance. A substantially higher proportion of specific mutations in a sample may suggest a bottleneck in viral entry or stochasticity in viral replication.

### Transcriptome changes at single-cell resolution

To understand transcriptional changes of the infected hAOs at a single-cell resolution, we employed two 10X Genomics single-cell RNA-seq experiments for uninfected and infected hAOs at 3 dpi (to a throughput of 21.3 Gb and 28.5 Gb, respectively). We captured 3,435 and 3,475 single-cells, respectively, with 15,703 UMIs and 3,414 detectable genes per cell on average. Using a total of 6,910 single-cells, we performed unsupervised clustering using UMIs of the human origin, and identified four distinct clusters (**Fig. 4a**): Cluster 1, mostly with uninfected alveolar cells (3,133 cells); Cluster 2, mostly with airway-like cells (361 cells; **Supplementary Discussion**); Cluster 3, alveolar cells with moderate levels of viral RNA transcripts (2,143 cells); and Cluster 4, cells disintegrating and likely close to cell death (n=1,273). The cells in the experiment of infected hAOs were mostly distributed in Cluster 3 (2,142 cells; 61.6% of infected hAOs) and Cluster 4 (1,154 cells; 33.2% of infected hAOs) (**Fig. 4b**). Viral transcripts were found in the vast majority of infected hAO cells (99.9%; 3,471 out of 3,475 infected hAOs), validating our previous observation that most cells in the infected hAOs harbor viral transcripts (**Fig. 2**). The number of viral transcripts, however, was not uniformly distributed in all infected hAO cells, but enriched in Cluster 4 cells. Infected cells in Cluster 4 exhibited 13.7 times more viral UMI counts than cells in Cluster 3, on average (**Fig. 4c;** 1,904 vs. 139 UMIs, respectively), despite cells in Cluster 4 containing relatively lower total UMI counts than ones in Cluster 3 (5,040 vs. 17,843 UMIs, respectively). When normalised with UMI counts for human genes, cells in Cluster 4 showed a >30 times higher viral RNA burden than cells in Cluster 3 (**Fig 4d**). Interestingly, the infected cells in Cluster 4 showed reduced expression of canonical hAT2 marker genes, including *SFTPB* (Surfactant Protein B) and *NKX2-1* (NK2 homeobox 1) (**Fig. 4e; Extended Data Fig. 6a**). Compared with infected cells in Cluster 3, expression levels of ISGs, such as *IFI44L* and *OAS3*, were also highly reduced. Instead, these cells showed transcriptional induction of apoptosis mediator, *GADD45B* (growth arrest and DNA-damage-inducible, beta) and anti-apoptotic *TNFAIP3* (tumor necrosis factor, alpha-induced protein 3), suggesting a catastrophic cellular pathway operating in a cell due to the extreme viral burdens.

**Figure 4.**
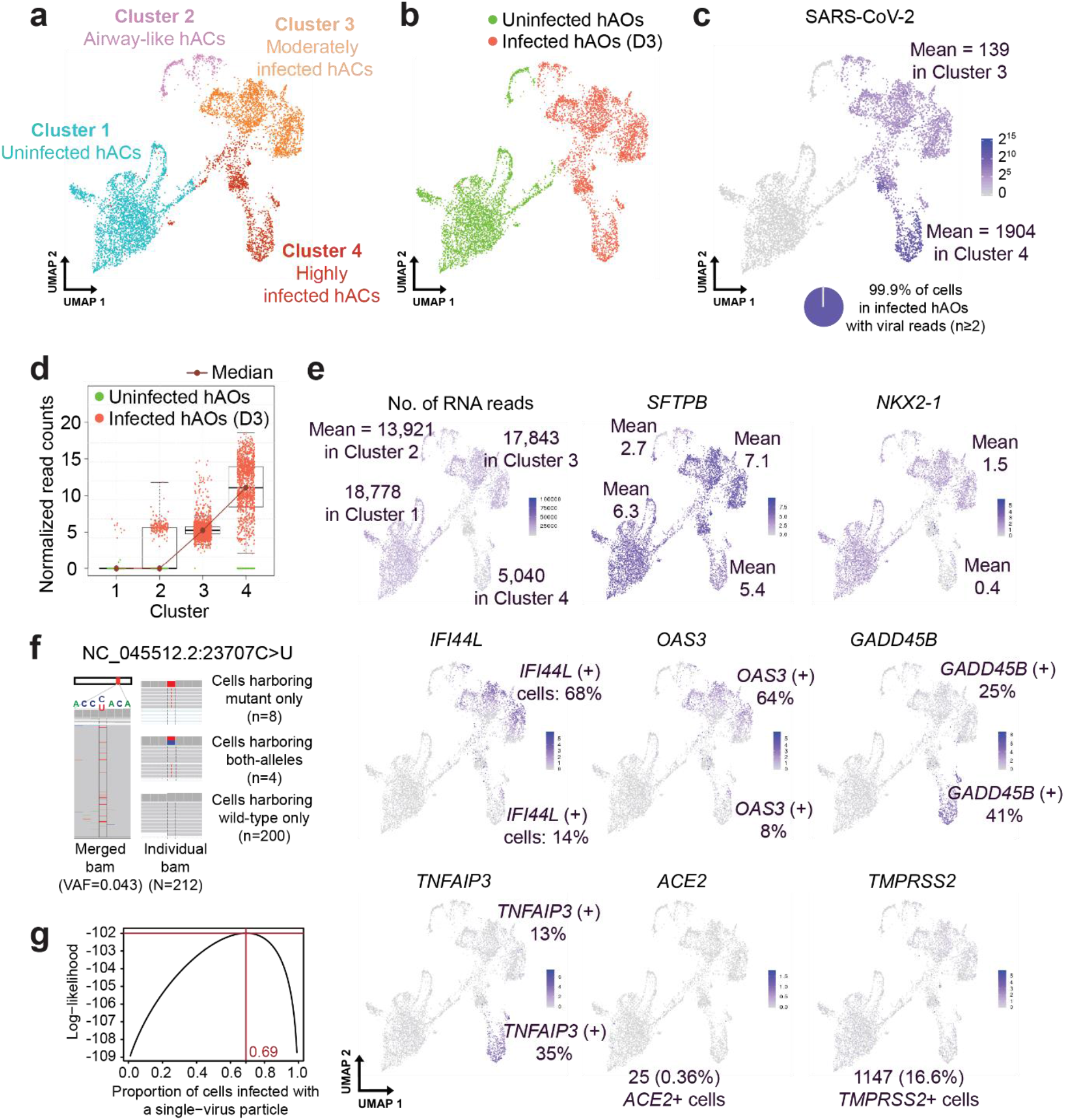
Single-cell transcriptome analysis of uninfected and infected hAOs. **a,** Unsupervised clustering of uninfected and infected human alveolar cells (3 dpi) in the UMAP plot. **b,** Experimental conditions of the dataset projected on the UMAP plot. **c,** Numbers of viral transcripts in each single-cell. Most cells harbor ≥2 viral reads. **d,** Normalized levels of SARS-CoV-2 viral read counts in each cluster. **e,** Expression of genes of interest in single-cells of each cluster. **f,** Distribution of single base substitution (variant allele fraction (VAF) = 4.3% in original viral particles) in each single cell. **g,** Maximum likelihood estimation for the proportion of cells infected with a single virus. This estimation indicates that infection by a single virus is more common in hAT2s rather than by multiple viruses.

Despite active protein expression (**Fig. 1f and 1g; Extended Data Fig. 1g**), we found 14 cells (0.4%) showing *ACE2* transcripts, 579 cells (16.7%) expressing *TMPRSS2* transcripts, and 4 cells (0.1%) co-expressing both in single-cell transcriptome sequencing (**Fig. 4e**). These proportions are low at face value, but are consistent with a previous observation^41^. Although the previous report also suggested that *ACE2* RNA expression can be stimulated as an infection-mediated response, particularly in human airway cells, such a trend was not observed in our dataset.

### Effective number of viral entry for infection of an alveolar cell

Finally, we statistically inferred the number of viral particles effectively entering each alveolar cell for infection. Although we incubated cells at an MOI of 1 on average, it is generally not known how many viral particles are necessary for effective infection of an alveolar cell. In an extreme scenario, one viral particle is sufficient. Alternatively, infection may be initiated with the entry of multiple viruses. We tracked the effective viral number of cellular entry using a mutation (NC_045512.2: 23,707C>U) as a viral barcode. From our sequencing, the mutation was estimated to be present at 4.3% VAF in the initial viral pool for innoculation. If the first scenario dominantly applies, the infected alveolar cells will be dichotomized, i.e., 95.7% cells with solely wild-type virus and the rest of the cells with solely mutant virus, exclusively. Alternatively, if multiple viruses are effectively entering a cell, a large proportion of alveolar cells harbouring detectable mutant virus will have intermediate VAF for the mutation. In our single-cell transcriptomes of infected hAOs, 212 cells were informative with the viral mutation, containing at least two independent transcripts for the mutant locus. The majority of the cells harbouring the mutant allele did not have the wild-type allele present (n=8 vs. 4 for cells showing solely mutant allele vs mixture of the quasispecies, respectively; **Fig. 4f**). In a more sophisticated statistical analysis, infection by single viral entry is estimated as >2 times more frequent than by multiple viral entry (69% vs. 31%, respectively; **Fig. 4g**). Our calculation indicates that a single viral particle is mainly responsible for SARS-CoV-2 infection in most alveolar cells, although multiple viral entry is also possible. It may also reflect the viral interference in SARS-CoV-2 alveolar infection.

## Discussion

In this study, we established conditions for optimised 3D long-term cultures of adult hAT2 cells, which provided an essential tool for studying initial intrinsic responses of SARS-CoV-2 infection. Single hAT2 cells were capable of self-constructing alveolus-like structures consisting of AT2 cell and differentiated AT1 cells. Mature hAT2 cells were maintained >10 months over multiple passages although self-renewal capacity and growth rate was reduced after 6 month in cultures. hAT1 cells were also lost from later culture, likely due to the persistent exposure to high WNT conditions allowing expansion of hAT2 cells over differentiation. Alteration of WNT activity in culture media (differentiation media) may enable to induce further AT1 cell differentiation.

hAOs showed remarkable phenotypic changes in the first few days after SARS-CoV-2 innoculation. Our single-cell transcriptome profiling identified two clusters of infected cells (Clusters 3 and 4), suggesting a distinct switch between cell states occurring during infection. If *in vivo* SARS-CoV-2 infection transforms hAT2 cells into Cluster 4-like cells showing the loss of hAT2 identity, the integrity and function of alveoli will be highly compromised. The route towards the quantum cellular change is of significant interest to be answered in future studies. To this end, we need more single-cell RNA sequencing from infected cells at earlier time points in combination with comprehensive high-resolution imaging.

The interferon response is the first line of host antiviral defense^40^. Contrary to a recent report^42^, we observed substantial ISGs in the alveolar cells induced by endogenously produced type I and III interferons. However, the induction of interferon responses was seen at 3 dpi in hAO models, 1-2 days later than the timing of viral amplification at 1 dpi. The timing of ISG induction may be earlier *in vivo* in concert with exogenous interferons from immune cells. For more physiological understanding, co-culturing SARS-CoV-2 infected hAO models with immune cells obtained from the same donor will be helpful.

In summary, our study highlights the power of feeder-free hAOs to elucidate the intrinsic responses of tissue damage including virus infection. Our data, including high-resolution electron microscopic images and the list of gene expression changes following infection, will be a great resource for the biomedical community to provide a deeper characterisation of SARS-CoV-2 infection specifically within adult hAT2 cells. We believe that our hAO models will enable more accurate and sophisticated analyses in the very near future, especially for studying the response of viral infection within vulnerable groups such as aged or diseased lungs, providing the opportunity to elucidate individual patient responses to viral infection. Furthermore, our models can be applied to other techniques, such as co-culture experiments with immune cells and robust *in vitro* screening of antiviral agents applicable to alveolar cells, in addition to being applicable for the study of the basic biology of alveolar cells as well as chronic disorders of the lung.

## Methods

### Human Tissues

For the establishment of human lung organoid models, human distal lung parenchymal tissues from deidentified lungs not required for transplantation were obtained from adult donors with no background lung pathologies from Papworth Hospital Research Tissue Bank (T02233), and Addenbrookes Hospital (Cambridge University NHS foundations trust) under the collaboration of Cambridge Biorepository for Translational Medicine (CBTM) project. Appropriate Human Tissue Act (HTA) guidance was followed; For organoids used for viral infection and following analysis, human lung tissue was obtained from patients undergoing lobectomy surgery at Seoul National University Hospital (SNUH) with written informed consent from approval of the ethical committee (approval no. C-1809-137-975). Organoids for infection studies were established from adjacent normal tissues in lung cancer patients or an idiopathic pulmonary fibrosis (IPF) patient.

### Virus particle preparation for infection

SARS-CoV-2 viral particles known as BetaCov/Korea/KCDC03/2020^6^,^31^ were used for the infection study. The patient (KCDC03) was diagnosed with COVID-19 on January 26, 2020, after traveling to Wuhan, China. The viral particles were prepared from Vero cells infected with MOI 0.01 and grown under DMEM (Sigma) with 2% FBS(Gibco), 1% P/S(Gibco) for 48 hours at 37°C 5% CO2. Media was centrifuged with 2500rpm for 25min, and supernatant without cell debris was stocked at −80°C with 4 × 10^6^ pfu / ml.

### Human lung tissue dissociation and flow cytometry

Distal lung parenchymal tissue was processed as soon as possible in order to minimize cell yield loss and maintain cell viability. Briefly, fresh tissue was washed in cold PBS and minced into small (1 mm) pieces with a scalpel, followed by further dissociation using pre-warmed digestion buffer containing 2U/ mL Dispase II (Sigma, Corning), 1 mg/mL Dispase/Collagenase (Sigma) and 0.1 mg/mL DNase I (Sigma) in PBS at 37 °C for 1 hr with agitation. Tissue cell suspensions were filtered through a 100 mM cell strainer into a 50 mL falcon tube to remove cell debris, and washed with 10 mL of DMEM (Gibco, Thermofisher). Cells were centrifuged at 350 g for 10 min, supernatant carefully aspirated, and cell pellet resuspended in 5 mL RBC lysis buffer for 5 min at room temperature (RT). The reaction was quenched using 5 mL of DMEM, and the entire 10 mL of cell suspension was transferred to a 15 mL falcon tube, followed by 10 min centrifugation at 350 g. Supernatant was removed, and the cell pellet was resuspended in 10% FBS in PBS (PF10 buffer) for counting. Cells were prepared for flow cytometry with primary antibodies CD31-APC (Biolegend, 303116), CD45-APC (Biolegend, 368512), EpCAM-FITC (Biolegend, 324204) and HTII-280-IgM (Terrace Biotech, TB-27AHT2-280) at 1:40 per 4 million cells for 30 min on ice. Following two washes with cold PF10 buffer and centrifugation at 350 g for 5 min, cells were stained with secondary PE goat anti-mouse IgM (eBioscience, 12-5790-81) for HTII-280 at 1:100. Stained cells were washed with PF10 buffer, and counted using a hemocytometer to assess dilution required for final volume. Cells were diluted at a concentration of 30 million cells/ mL and filtered through a 35 μM cell strainer into polypropylene FACS tubes. Cell sorting was performed on an Aria III fusion (BD Biosciences) using a 100 μM nozzle, and data were analysed with FlowJo software (Tree Star, Inc.).

### *In vitro* organoid culture and passage

Freshly isolated HTII-280^+^ and HTII-280^−^ cells derived from CD31−CD45−EpCAM^+^ cells of human distal lungs were resuspended in Advanced DMEM/F12 (Thermofisher) base medium supplemented with 10 mM Hepes (Gibco), 1 U/mL Penicillin/Streptomycin (Gibco), 1 mM *N*-Acetylcysteine (Sigma), and 10 mM Nicotinamide (Sigma). Growth factor-reduced (GFR-) Matrigel (Corning) was added to the cell suspension at a ratio of 1:1, and 100 μL of suspension was added to a 24-well transwell insert with a 0.4 μM pore (Corning) so that there were approximately 10 × 10^3^ cells per insert. GFR-Matrigel was allowed to solidify for 1 hr at 37 °C, after which 500 μL of pre-warmed alveolar media (base media supplemented with 1 × B27 (ThermoFisher), 10% R-SPONDIN-1 (Cambridge Stem Cell Institute tissue culture core facility, manually produced), 50 ng/ml human EGF (Peprotech), 100 ng/ml human FGF7/KGF (Peprotech),100 ng/ml human FGF10 (Peprotech), 100 ng/ml NOGGIN (Peprotech),10 μM SB431542 (Tocris) and 3 μM CHIR99021 (Tocris)) was added to each lower chamber. For assessment of the effect of Wnt activity on hAT2 culture ability, primary cultures were also established without CHIR99021. Cultures were maintained under standard cell culture conditions (37 °C, 5% CO_2_), with media changes every 2-3 days. Y-27632 (10 μM, Sigma) was added for the first 48 hr of culture to promote cell survival. To avoid the growth of fungal and bacterial infection, 250 ng/mL Amphotericin B and 50 μg/mL gentamicin were added to culture medium for 5 days. For culture in 48-well plates, 5 × 10^3^ cells were resuspended in 100% GFR-Matrigel, and allowed to solidify in a 20 μL droplet per well at 37 °C for 20 min, followed by submersion in 250 μL of pre-warmed medium. Organoids larger than 45 μM in size were counted at day 14 of culture to assess organoid forming efficiencies, and were either fixed and stained at day 21 for analysis, or enzymatically dissociated into single cells for further culture without sorting. Organoid lines were passaged at different days depending on size, with culture days varying from 21-35 days. For passaging, Matrigel was disrupted by incubation with Dispase (Sigma) at 37 °C for 45 min, followed by single cell-dissociation through addition of trypLE (Gibco) for 5 min at 37 °C. The reaction was quenched with base medium, and cells were centrifuged at 350 × g for 5 min. Cells were resuspended in fresh GFR-Matrigel at a ratio of 5 × 10^3^ (48-well plates) or 10 × 10^3^ (24-well transwell inserts) cells as before. HTII-280^−^ cells were cultured as with HTII-280^+^, although with a few minor differences. Bronchial organoids (hBOs) were passaged every 21-28 days due to accelerated growth compared with alveolar organoids (hAOs), and were cultured in previously reported medium conditions^30^ with the following concentration/factor edits; 100 ng/ml human FGF10, 10% R-SPONDIN-1, 10 μM SB431542 (instead of A83-01).

### Virus infection to organoids and Vero cells

For hAO and hBO, cells were recovered from Matrigel with a Recovery solution (Corning). Organoids were sheared with 1000 p pipette tips or incubated with Accutase (Stem cell technology) at 37 °C for 5 minutes. Organoids were resuspended with each organoid’s media and were infected with virus multiplicity of infection (MOI) of 1 for 2 hrs at 37 °C 5% CO_2_. After virus infection, organoids were washed twice with Advanced DMEM/F12 with 1 U/ml Penicillin/Streptomycin, 10 mM Hepes, and 1% Glutamax (v/v) (hereafter referred to as ADF+++) and embedded with 50 μl of GFR-Matrigel (Corning) in 24-well plate (TPP). At least each well contained 10,000 cells. Media and cells resuspended in ADF+++ with Matrigel were taken at indicated time points. For viral particle release inside cells, cells were lysed by freezing at −80 °C and thawing. Live virus titers were determined by plaque assay and viral RNA titer was calculated using qPCR. For Vero cell infection, a virus with MOI 1 was poured on Vero cells directly, and cells with viruses were incubated for 1 hr at 37 °C 5% CO_2_. After infection, infection media was removed and cells were incubated in normal media in 37 °C 5% CO_2_ condition. All work was performed in a Class II Biosafety Cabinet under BSL-3 conditions at Korea Center for Disease Control (KCDC).

### Immunofluorescence staining of paraffin-embedded organoids

Organoids were fixed and embedded in a paraffin block^43^. Pre-cut 7 μM paraffin sections were de-waxed and rehydrated (sequential immersion in xylene, 100% EtOH, 90% EtOH, 75% EtOH, distilled water) and either stained with hematoxylin and eosin (H&E) or immunostained. For antigen retrieval, slides were submerged into pre-heated citrate antigen retrieval buffer (10 mM sodium citrate, pH 6.0) and allowed to boil for 15 min. Slides were cooled in a buffer for 20 min, washed in running water for 3 min, and permeabilised with 0.3% Triton-X in PBS for 15 min. Following permeabilisation, organoid sections were blocked for 1 hr in 5% normal donkey serum in PBS at RT, and incubated with primary antibody mixes overnight at 4 °C at the following dilutions; rabbit pro-SFTPC (1:500, Millipore, Ab3786), mouse anti-HTII-280 (1:500, Terrace Biotech, TB-27AHT2-280), rat anti-SCGB1A1 (1:200, R&D systems, MAB4218), rabbit anti-KRT5 (1:500, Biolegend, 905501), mouse anti-ABCA3 (1:300, Seven Hills Bioreagents, WRAB-ABCA3), and mouse anti-TP63 (1:500, Abcam, ab735), rabbit anti-HOPX (1:200, Santa Cruz Biotechnologies, sc-30216), and sheep anti-PDPN (1:200, R&D, AF3670). Antibodies were removed with three PBS washes, and samples were incubated with Alexa Fluor-couple secondary antibodies (1:1000, Jackson Laboratory) for 1 hr at RT. Following PBS washes, nuclei were stained with DAPI for 5 min, slides mounted with Rapiclear (Sunjin lab), and sealed with clear nail polish. For Lysotracker™ staining of lysosomes, live organoids in 48-well plates were incubated *in situ* with 50 ng/μL of Lysotracker™ (Invitrogen, L12492), diluted in pre-warmed expansion medium, for 1 hr at 37 °C. Lysotracker was removed, and organoid/matrigel suspension was carefully washed for 5 min in PBS, followed by addition of fresh, pre-warmed expansion medium. Cells were protected from light and imaged immediately using an EVOS cell imaging system.

### Immunofluorescence staining of infected organoid with cryosection

Organoids were fixed in 4% paraformaldehyde (PFA) for 3 hrs at ice, and then dehydrated in PBS with 30% sucrose (v/v) (Sigma). Organoids were embedded with optimal cutting temperature (OCT) compound (Leica) and cut with 10 μM. Organoid section was blocked with 5% normal donkey serum in PBS 1% triton-X (Sigma). Sections were incubated with primary antibodies overnight at 4 °C, washed three times with PBS. Organoids were incubated host matched Alexa Fluor-couple secondary antibodies (Jackson Laboratory) with PBS for 1.5 hr at RT. Following DAPI incubation, slides were mounted. Antibodies were E-cadherin (R&D, AF748, 1:300), NP (Sino, 40143-MM05, 1:200), Spike S1 protein (Sino, 10030005, 1:200), ACE2 (abcam, ab15348, 1:400), TMPRSS2 (abcam, EPR3861,1:400)

### Transmission electron microscopy

Human alveolar organoids were fixed with 2.5 % glutaraldehyde in 0.1 M PBS overnight at 4 °C ^44^. Organoids were washed with PBS and post-fixed with 2% osmium tetroxide for 1.5 hr. The fixed sample was dehydrated in graded ethanol, substituted with propylene oxide, and finally embedded in EMbed-812 resin (EMS). Polymerization was performed at 60 °C for 24 hrs. Ultrathin (100 nm) sections were prepared using an ultramicrotome (Leica, EM UC7). Images were captured with a transmission electron microscope (FEI Tecnai G2 spirit TWIN, eagle 4K CCD camera) at 120kV acceleration voltage. All work was carried out in the EM & Histology Core Facility, at the BioMedical Research Center, KAIST.

### RNA purification

For viral RNA extraction, cells were resuspended with Matrigel with media and frozen at −80 °C and then thawed. Thawed solution was processed through the QIAamp Viral RNA Mini Kit. For mRNA extraction, cells were recovered from Matrigel by a Recovery solution (Corning) and then centrifuged at 300 g for 5 min at 4 °C. Cell pellets were lysed, and RNA was extracted through RNeasy Plus Mini Kit. For viral RNA extraction from media, 140 μl of media was processed through QIAamp Viral RNA Mini Kit.

### Assessment of lung lineage transcripts by qRT-PCR

Freshly sorted HTII-280^+^ and HTII-280^−^ cells were lysed with TRIzol, and RNA was extracted. RNA was reverse transcribed using SuperScript IV (Thermo Fisher Scientific), and were assessed using the following Taqman probes; SFTPC (Hs00951326_g1), TP63 (Hs01114115_m1), SCGB1A1 (Hs00171092_m1).

### Viral RNA copy number calculation with qRT-PCR

Viral RNA samples were reverse-transcribed using SuperScript IV (Thermo Fisher Scientific). Viral N3 gene was targeted for qRT-PCR. Nucleotide sequences of the probes as below (CDC).

2019-nCoV_N3-F: 5’ GGGAGCCTTGAATACACCAAAA 3’
2019-nCoV_N3-R: 5’ TGTAGCACGATTGCAGCATTG 3’

Samples were prepared triplicate from solution. Viral RNA copy number was calculated by comparing the standard curve of virus N3 gene. For generating the standard curve, the positive viral RNA template was amplified with CDC designed N3 gene primers and cloned into pGEM-T Easy vector (Promega, USA). The resultant plasmid DNA was linearized with PstI restriction enzyme and purified with a QIAquick PCR Purification Kit (QIAGEN, Germany). Purified template was in vitro transcribed by RiboMAX™ Large Scale RNA Production System with T7 RNA polymerase (Promega, USA). RNA transcript was further purified with the NuceloSpin RNA Mini kit (MACHEREY-NAGEL, Germany) and quantified with spectrophotometer at 260 nm. *In vitro* transcribed RNA was serially diluted and reverse transcribed for the quantitative real time PCR^45^.

### Live virus titer calculation with plaque assay

Matrigel was sheared with the organoid media and frozen at −80 °C once. Thaw the solution and dilute by scale of 10. Each well containing Vero cells in 12 wells were infected with the diluted solution respectively at 37 °C, 5% CO_2_ for 1 hr. After infection, remove infection media and wash the Vero cells with PBS two times, mixed agar and Modified Eagle’s Medium (Thermofisher) were poured on each well. When agar mixture was hardened, fix each well with 4% PFA for 3 days, and stain with crystal violet (Sigma). When there are individual spots, the original solution’s viral titer was calculated.

### Bulk RNA sequencing and data processing

Extracted cellular RNA was processed through Truseq Stranded Total RNA Gold kit, and cDNA library was sequenced 2 × 100 bp using Hiseq 2500. Fastq file was aligned to GRCh38 with virus sequence (NC 045512.2 from NCBI) using STAR^46^ and normalized RNA expression was calculated using RSEM^47^. Differentially expressed genes are found from DEseq2^48^. We obtained enriched gene sets using in-house scripts. For mutation calling, we used Strelka2^49^, Varscan2^50^, and Samtools^51^, and then manually checked the position through IGV^52^.

### Single-cell RNA sequencing and data processing

Fastq file was aligned and each UMI count was calculated using Cell Ranger software provided by the manufacturer (10X Genomics). Cells with mitochondria RNA percent < 25%, total RNA number > 200 subsets were used for downstream analysis.

Starting from the 10X gene counts, we have normalized data as follows. The 10X data includes SARS-CoV-2 genome as an extra gene besides 19,941 human genes. For uninfected cells, such SARS-CoV-2 gene would have zero read count, while for infected cells, the gene could account for a large portion of total reads. One of the goals of normalisation is to remove technical difference such as sequencing coverage before comparing gene expression levels. As our interest is to compare expression levels between uninfected and infected cells on human genes, we have applied a normalisation method (‘scater R package’s ‘logNormCounts’ function) to human genes as a whole set. The method calculated a scaling factor for each cell based on total human gene count, then scaled all genes before taking log-transformation. Using the same scaling factor learned during human gene normalisation, SARS-CoV-2 count was also normalized. For clustering, we combined single-cell data from infected and uninfected hAOs. Unsupervised clustering was performed using a shared nearest neighbor (SNN) based clustering algorithm in Seurat^53^. Contaminated cells (< 3%) were discarded. In house R scripts were used for more downstream analyses.

### Statistical inference on the effective number of viral entry

To assess whether alveolar cells tend to be infected by a single viral particle or multiple particles, we employed a likelihood approach. As a proof-of-concept, we assumed only two scenarios exist, one supporting a single viral entry and the other supporting double viral entry, then aimed to estimate the proportion of cells with a single viral particle (*w*). Out data consists of the observed reference 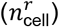 and variant 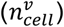 read counts for each of the 547 reporting at least one read at the mutation site of NC_045512.2:23,707. Assuming a sequencing error rate (∊) of 0.1%, which will cover any Illumina sequencing errors or misalignment, the likelihood of data given the weight supporting a single virus scenario (*w*) was computed as follows.

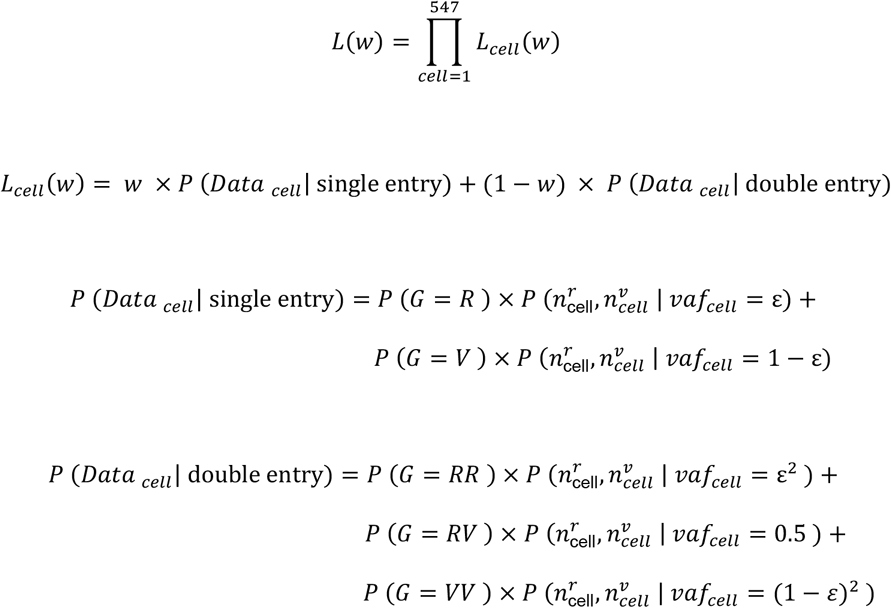

*G* stands for the viral genotype within each cell, and *R* and *V* denote for the wildtype and mutant alleles in the genotype, respectively. *vaf*_*cell*_ refers to the true mutant allele frequency within an infected cell given the viral genotype after accounting for the sequencing error rate.

For both single- or double-entry scenarios, the prior for each viral genotype was computed using the mutant allele frequency observed from the whole cell population (*vaf*_*pop.*_; 4.3%=42 variant reads out of 986 reads). For example, the prior of having the genotype with two mutant alleles in a double-entry scenario is computed as follows:

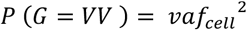

Within a cell, the probability of observing the number of reference and variant reads given the mutant allele frequency, 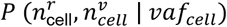, was computed using binomial distribution.

## Supplementary discussion

### Basal-like cells and hAT2 enrichment in single cell RNA sequencing

We observed a distinct transcriptional feature of cells in Cluster 2 which express lower levels of canonical hAT2 marker genes, including *SFTPC* but detectable levels of airway marker genes, including *SOX2*, *TP63*, *KRT5*, and *KRT17* (**Extended Data Fig. 6b**). These expression patterns were not affected by virus infection. hAT2 cells expressing airway markers such as SOX2 were seen in chronic lung diseases such as lung cancer and idiopathic pulmonary fibrosis (IPF)^54,55^, representing pathologic phenotypes of alveolar bronchiolization. A recent study also suggested the potential transition of hAT2 cells to KRT5^+^ basal-like cells in the context of IPF^27^. Given the fact that the hAOs used for our single-cell RNA sequencing study were derived from hAT2 cells isolated from adjacent normal counterparts of lung cancer and/or IPF, it is likely that this transcriptional feature reflects the cellular status of original tissues rather than virus-associated phenotype. This finding suggests that our hAO models maintain the pathophysiologic features of original tissues although we used apparently normal background regions for our hAO establishments. Further long-term tracing of changes in cellular identities and states in response to virus infection in hAO cells will be of significant interest to understand the progression of pathologic features and reparative mechanisms for developing therapeutic interventions.

Furthermore, from our scRNAseq analysis, most captured cells were hAT2 cells. It is likely that this might result from the enrichment of hAT2 cells in our hAOs (P2) and the nature of fragile hAT1 cells during the procedure of single-cell preparation for scRNAseq.

### Supplementary Tables

**Supplementary Table 1**. RNA expression levels (TPM) of all genes in seven human alveolar organoid samples.

**Supplementary Table 2**. RNA expression levels (TPM) of all genes in seven human bronchial (airway) organoid samples.

**Supplementary Table 3**. Twenty single base substitutions of SARS-CoV-2 which were detected in infected organoids.

## Acknowledgements

We thank Jinwook Choi (Wellcome-MRC Cambridge Stem Cell Institute), Yong Man Han, Eui-Cheol Shin, Heung Kyu Lee, Su-Hyung Park, Jeong Seok Lee, Ryul Kim, Myungsuk Choi (KAIST), Jung-Ki Yoon (Seoul National University Hospital), Yoon Ho Kim (Seoul National University Cancer Research Institute), and Jong-Yeon Shin (Macrogen Inc.) for valuable discussion, productive comments, and technical help; Irina Pshenichnaya (Histology), Peter Humphreys (Imaging), Simon McCallum (Flow cytometry, Cambridge NIHR BRC Cell Phenotyping Hub), and Cambridge Stem Cell Institute core facilities for technical assistance. This work was supported by Suh Kyungbae Foundation (SUHF-18010082 to Y.S.J.); National Research Foundation of Korea (for Brain Pool Program NRF-2019H1D3A2A02061168 to Y.S.J and S.Y.K; Leading Researcher Program NRF-2020R1A3B2078973 to Y.S.J) and Institute for Basic Science (IBS-R025-D1, G.Y.K) funded by Ministry of Science and ICT of Korea. J.-H.L was supported by Wellcome and the Royal Society (107633/Z/15/Z) and European Research Council Starting Grant (679411). K.V.E was supported by the Biotechnology and Biological Sciences Research Council Industrial CASE (BBSRC iCASE) studentships (BB/R505328/1).

## Author contributions

Conceptualization, Y.S.J. and J.-H.L.; hAO development, K.V.E., J.-H.L.; hAO establishments and characterisation, K.V.E., J.Y., T.K., J.-H.L.; Specimen preparation; K.J.N, K.M., K.S., Y.T.K., K.M., K.S.P.; Infection, J.K., J.Y., T.K., Y.-I.J., B.-S.C; Confocal microscope, K.V.E., S.P.H., G.Y.K.; Electron microscope, Y.H., T.K., H.M.K.; Transcriptome analysis, J.Y., T.K., K.Y., S.Y.K., Y.S.J.; Statistics, S.Y.K.; Writing, J.Y., T.K., K.V.E., T.B., S.-H.K., G.Y.K., Y.S.J., J.-H.L.; Manuscript finalization, all authors; Supervision Y.S.J. and J.-H.L.

## Declaration of interests

The authors declare no competing interests.

## Materials availability

All unique organoids generated in this study are available from Young Seok Ju or Joo-Hyeon Lee with a completed Materials Transfer Agreement.

## Data availability

Bulk RNA and single cell RNA sequencing datasets will be uploaded on the European Genome-Phenome Archive (EGA). Accession ID is not assigned yet.

---

**Extended Data Figure 1.**
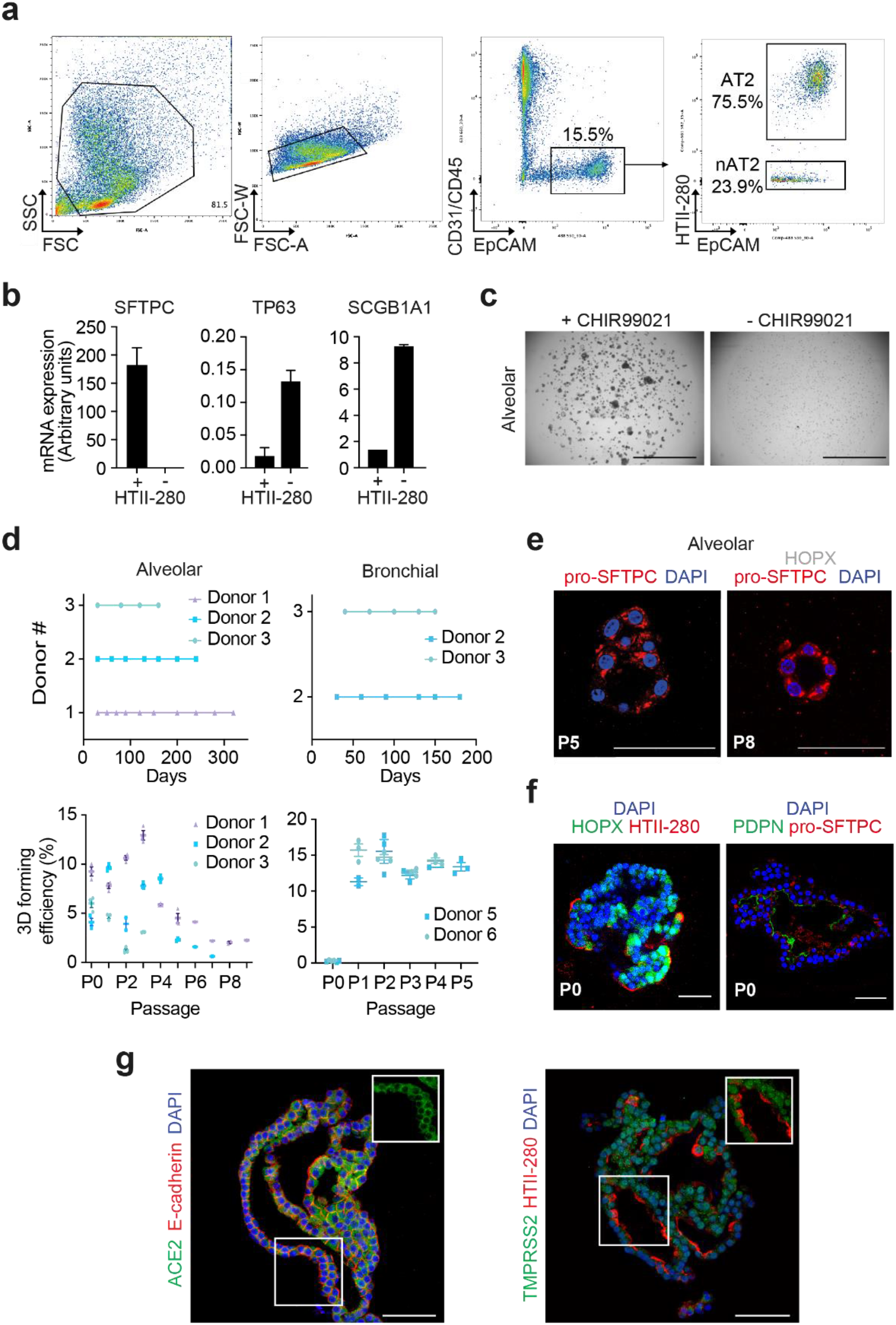
Establishment and characterisation of three-dimensional culture of human alveolar type 2 cells. **a,** Representative flow cytometry analysis for isolation of hAT2 cells (CD31^−^CD45^−^EPCAM^+^HTII-280^+^) and non-hAT2 cells (nAT2, CD31^−^CD45^−^EPCAM^+^HTII-280^−^) from human distal lung parenchymal tissues. **b,** Quantitative PCR (qPCR) analysis of isolated primary human epithelial lung cells in (a) for lung lineage marker genes revealed that HTII-280^+^ cells expressed much higher level of the AT2 cell marker SFTPC, while HTII-280^−^ cells expressed virtually no SFTPC and instead expressed higher levels of the airway markers TP63 and SCGB1A1. Data is the mean ± SEM of two technical replicates, and is expressed as arbitrary mRNA expression. **c,** Representative bright-field images of organoids derived from hAT2s in complete medium (left) and withdrawal of CHIR99021 (right). Scale bar, 2000 μm. **d,** Top; Organoid cultures for alveolar or bronchial cells from different donors were passaged at various time points depending on growth. Each point represents a single passage. Bottom; Statistical quantification of organoid forming efficiency at day 14-28 up to 9 passages (alveolar) and 5 passages (bronchial). Each individual dot represents one technical replicate, and data are presented as mean ± SEM for each individual donor sample. N=3 for hAOs; N=2 for hBOs. **e,** Organoids continue to express pro-SFTPC (red) throughout culture (passage 5, left; passage 8, right), even following 9 months in culture, but do not exhibit expression of the AT1 marker HOPX (white) during later passages. DAPI (blue). Scale bar, 50 μM. **f,** Primary hAOs derived from hAT2 cells demonstrate expression of the AT1 markers HOPX (green, left) and PDPN (green, right) and the AT2 markers HTII-280 (red, left) and pro-SFTPC (red, right panel). DAPI (blue). Scale bar, 50 μm. **g,** Immunofluorescent staining of ACE2 and TMPRSS2 (green) is consistent in hAOs derived from different donors. Scale bar, 50 μm.

**Extended Data Figure 2.**
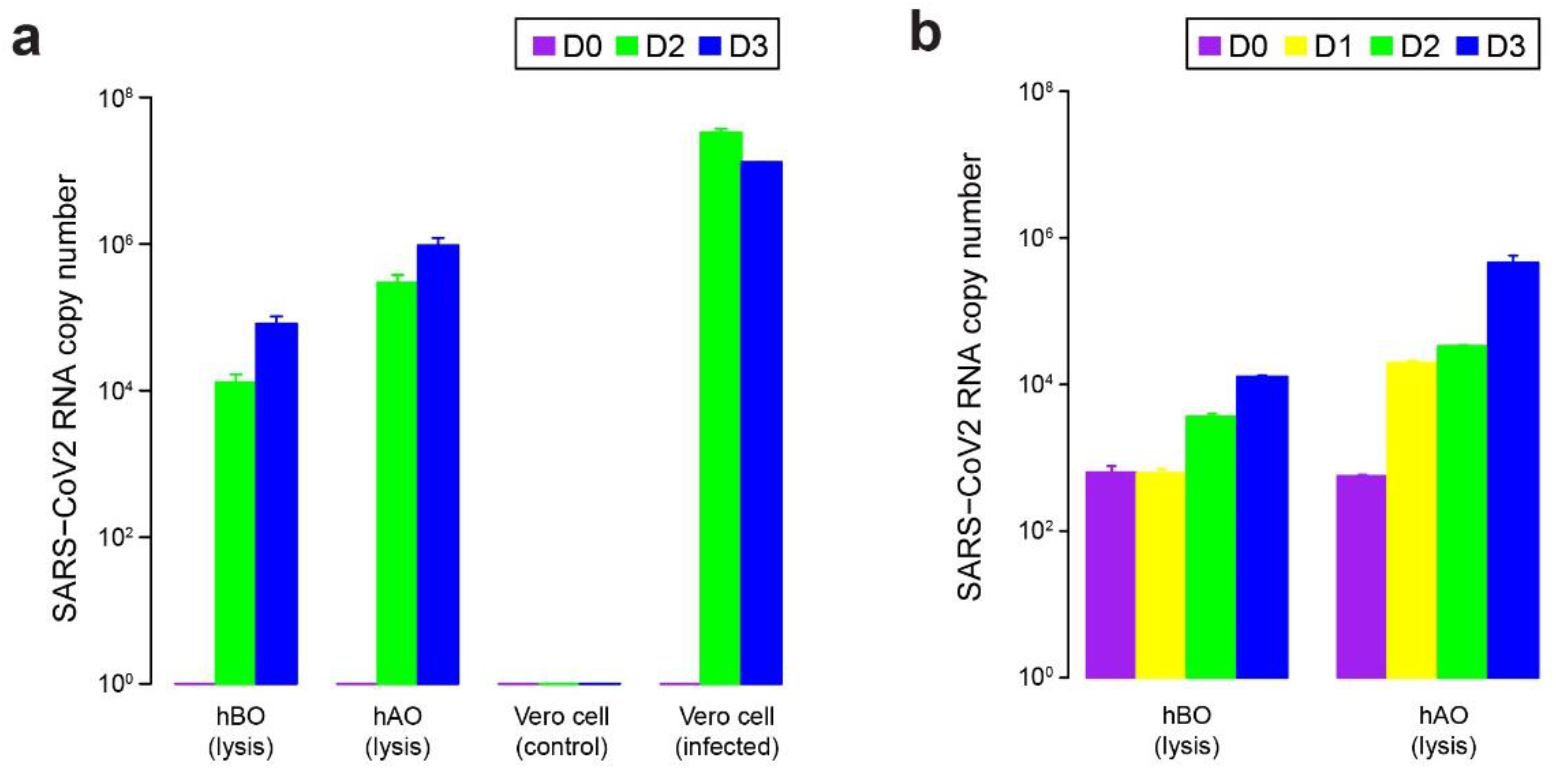
Repeated experiments of SARS-CoV-2 infectivity in three-dimensional culture of human alveolar type 2 cells. **a,** qPCR analysis for measuring the viral RNA levels in hBOs, hAOs, and vero cells (Replicate #2). SARS-CoV-2 infectivity is much higher (<100 times) in hAOs rather than hBOs. Error bars represent SEM. n=2. **b,** qPCR analysis for measuring the viral RNA levels in hBOs and hAOs. (Replicate #3). Error bars represent SEM. n=3

**Extended Data Figure 3.**
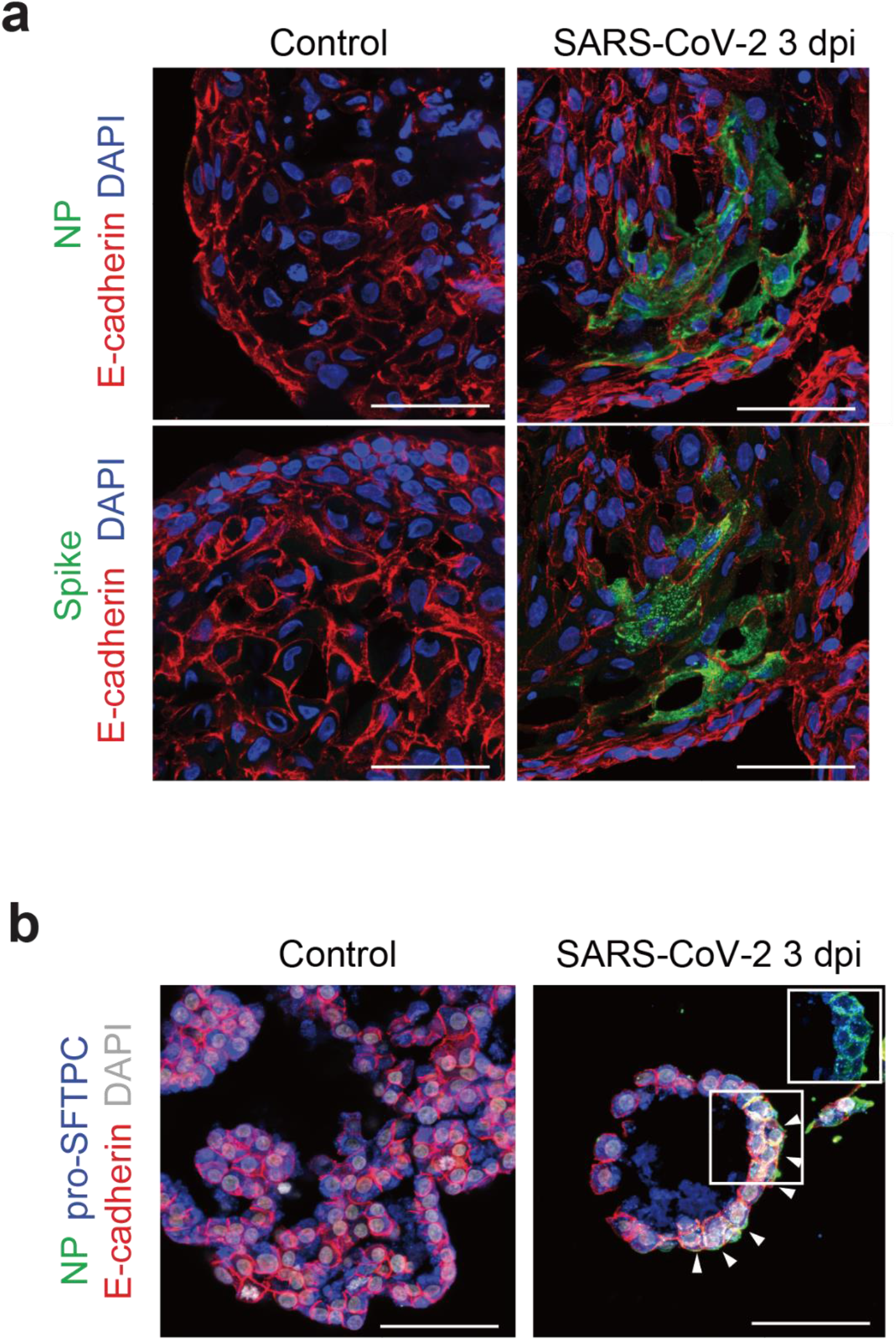
Immunofluorescent images of human alveolar type 2 organoids at 3 days after SARS-CoV-2 infection. **a,** In a portion of the hAO at 3 dpi, SARS-CoV-2 infection is identified by viral nucleoprotein (NP) and spike protein. Scale bar, 50 μm. **b,** SARS-CoV-2 infected hAOs still express the AT2 cell marker pro-SFTPC. Scale bar, 50 μm.

**Extended Data Figure 4.**
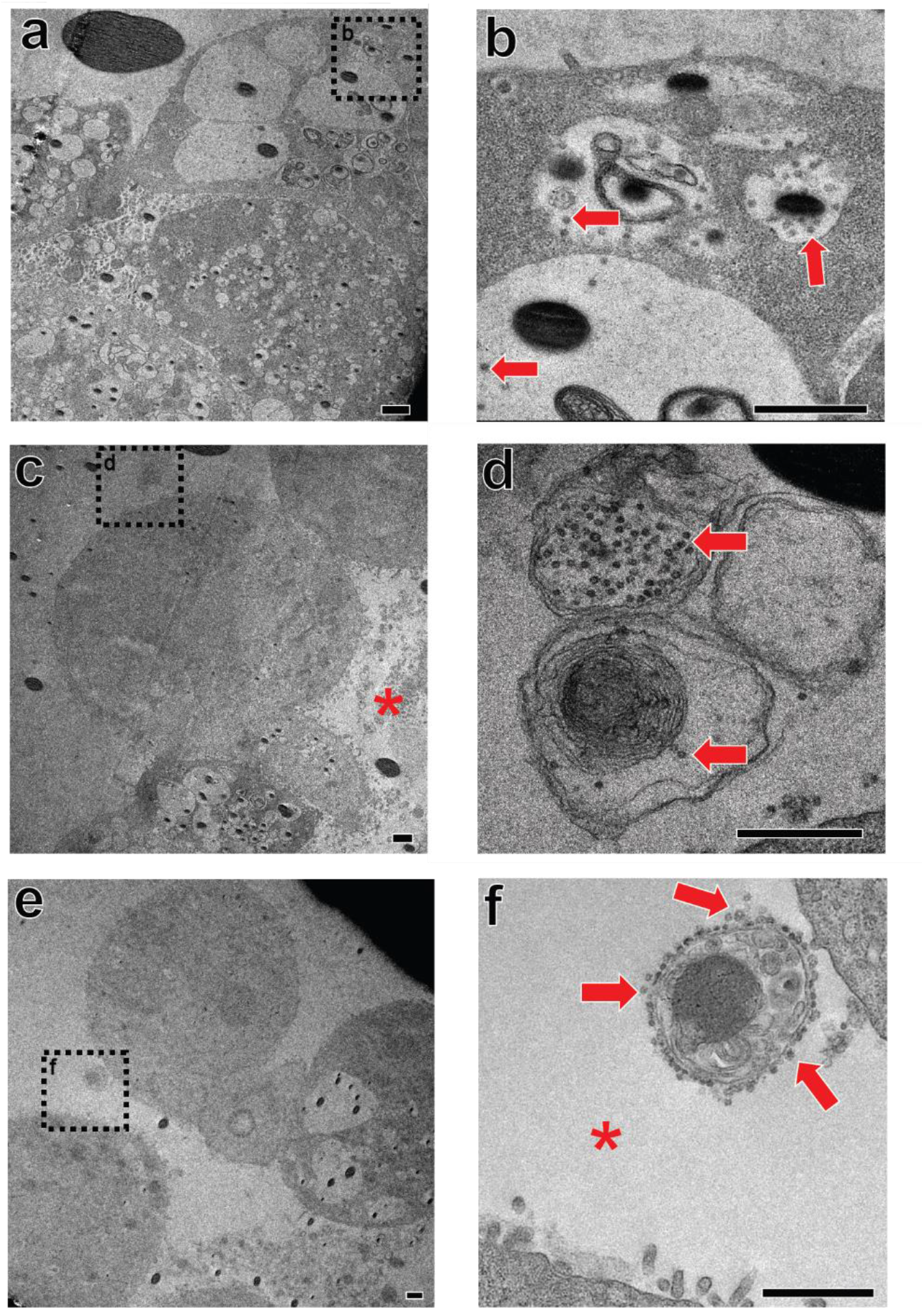
Electron microscopy analysis of human alveolar type 2 cells infected by SARS-CoV-2. **a,** A low magnification image of an infected hAO. **b,** Viral particles (red arrow) in a large vacuole. The density of viral particles is lower than that of small vesicle structures. **c,** Another low magnification image of an infected hAO. Alveolar space is annotated by red asterisk. **d,** Viral particles are encapsulated in the exocytic vesicles (red arrow). **e,** A low magnification of hAT2 cells. **f,** Viral particles are attached to the outgoing vesicle (red arrow). Viruses are secreted in different ways. Scale bar, 1 μm.

**Extended Data Figure 5.**
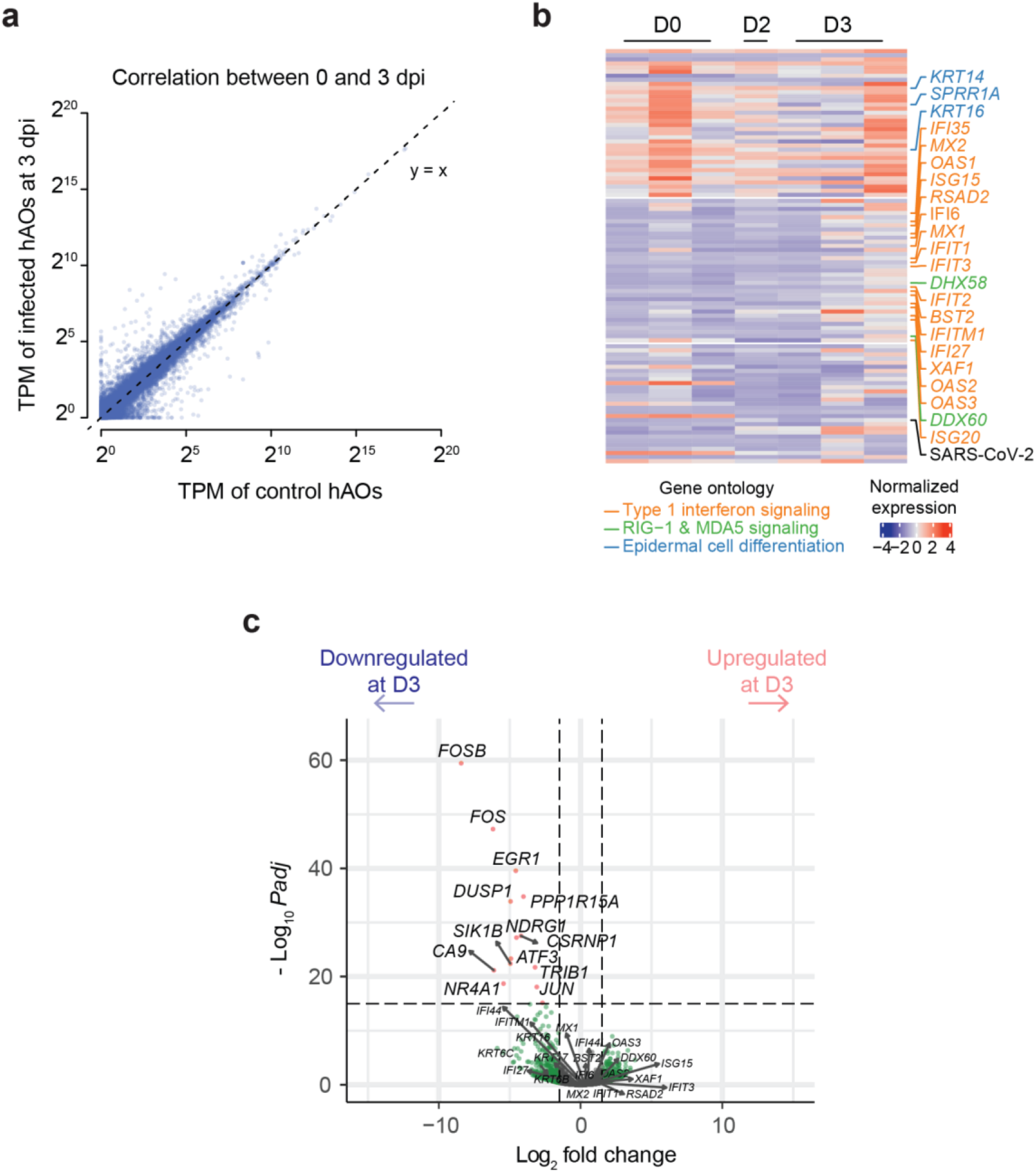
Transcriptome changes of the infected human lung organoids. **a,** Correlation of TPM level of each gene in hAOs at 0 and 3 dpi. Each gene is depicted as a blue dot. **b,** Heatmap of the most variable 100 genes derived from hAOs (Figure 5A, Table S1) in SARS-CoV-2 infected hBOs. Normalisation of gene expression was calculated using all transcriptome data from 14 samples of hAOs and hBOs. **c,** Volcano plot for differentially expressed genes in hBOs at 0 and 3 dpi. No significant changes were shown in ISGs.

**Extended Data Figure 6.**
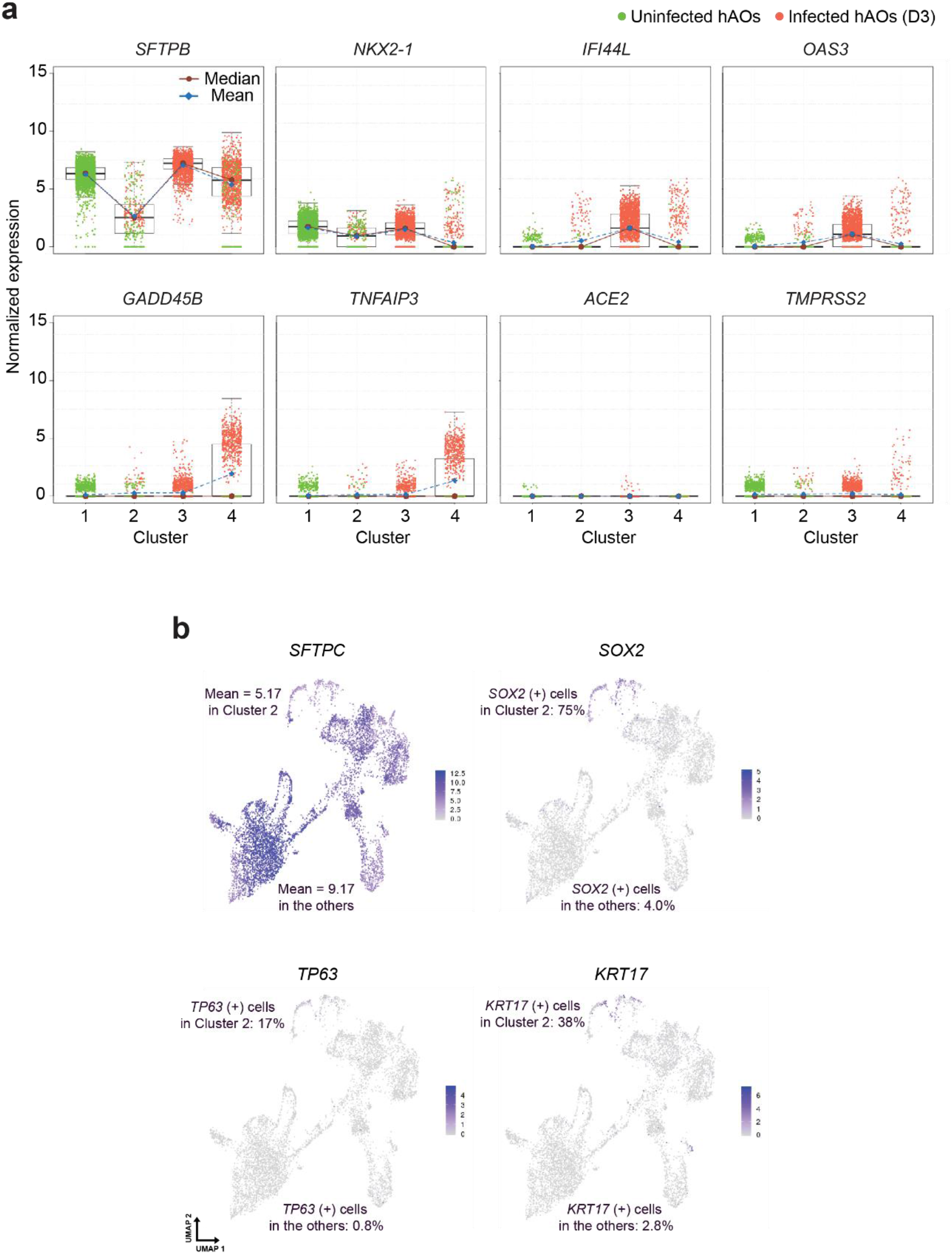
Single-cell transcriptome profiling of infected hAOs. **a,** Box plots showing normalised expression of interested genes in each cluster. Center line, median; box limits, upper and lower quartiles; whiskers, 1.5x interquartile range. **b,** In Cluster 2, expression level of *SFTPC*, one of hAT2 cell marker genes is reduced. By contrast, some airway marker genes such as *SOX2*, *TP63*, and *KRT17* were expressed.

## Notes

### Competing Interest Statement

The authors have declared no competing interest.

